# TK216 targets microtubules in Ewing sarcoma cells

**DOI:** 10.1101/2021.11.19.469182

**Authors:** Juan Manuel Povedano, Vicky Li, Katherine E. Lake, Xin Bai, Rameshu Rallabandi, Jiwoong Kim, Yang Xie, Jef K. De Brabander, David G. McFadden

## Abstract

Ewing sarcoma (EWS) is a pediatric malignancy driven by the EWSR1-FLI1 fusion protein formed by the chromosomal translocation t(11;22). The small molecule TK216 was developed as a first-in-class direct EWSR1-FLI1 inhibitor and is in phase II clinical trials in combination with vincristine for EWS patients. However, TK216 exhibits anti-cancer activity against cancer cell lines and xenografts that do not express EWSR1-FLI1, and the mechanism underlying cytotoxicity remains unresolved. We apply a forward genetics screening platform utilizing engineered hypermutation in EWS cell lines and identify recurrent mutations in *TUBA1B*, encoding α-tubulin, that prove sufficient to drive resistance to TK216. Using reconstituted microtubule (MT) polymerization in vitro and cell-based chemical probe competition assays, we demonstrate that TK216 acts as an MT destabilizing agent. This work defines the mechanism of cytotoxicity of TK216, explains the synergy observed with vincristine, and calls for a reexamination of ongoing clinical trials with TK216.

## MAIN TEXT

EWS is defined by chromosomal translocations that lead to the expression of oncogenic fusion proteins of the EWSR1-FLI1 family. The EWSR1-FLI1 family of proteins is formed by the fusion of protein sequences of low complexity from EWSR1, FUS or TAF15 to the DNA-binding domain of an E26 transformation-specific (ETS) family transcription factor, most frequently FLI1 or ERG^1^. The EWSR1-FLI1 protein lacks known enzymatic activity or defined small molecule binding pockets; therefore, rational therapeutic targeting of the protein represents a major challenge.

YK-4-279 is the first small molecule reported to directly target EWSR1-FLI1. It was identified as a small molecule capable of disrupting an interaction between EWSR1-FLI1 and RNA helicase A (encoded by DHX9). YK-4-279 induced apoptotic cell death in EWS cell lines and suppressed growth of EWS xenografts^2^. YK-4-279 was subsequently shown to induce G_2_-M cell cycle arrest and apoptosis in synergy with the MT destabilizing agent vincristine^3^. TK216, a clinical derivative of YK-4-279, has entered phase II clinical trials in EWS patients as monotherapy and in combination with vincristine, and a subset of patients exhibits promising responses^4^.

Since the original identification of YK-4-279 as an inhibitor of EWSR1-FLI1, the molecule has been shown to suppress growth of a variety of cancer cell lines not driven by EWSR1-FLI1, including prostate cancer, neuroblastoma, lymphoma, melanoma and thyroid cancer^5–9^. In addition, genetic suppression of EWSR1-FLI1 induced cell cycle arrest at a different checkpoint: the G_1_-S transition^10^. These findings have raised questions regarding the mechanism underlying cytotoxicity induced by YK-4-279, which remains unresolved.

We previously reported that engineered DNA mismatch repair (MMR) deficiency induces hypermutation in cancer cell lines that facilitates the emergence of compound resistant alleles^11^. These mutations can reveal the direct protein targets of cytotoxic small molecules^11–14^. We sought to uncover the mechanism of action of TK216-induced cytotoxicity using this unbiased forward genetics platform. We performed forward genetic screening with TK216 using MMR-deficient A673 EWS cells at three concentrations flanking the IC_100_^1wk^ (Fig. 1A, Methods). Six compound-resistant clones (TK216 A-F) emerged following TK216 selections. We confirmed resistance to TK216 in all six clones (range from 1.98- to 2.74-fold compared to parental MMR-deficient A673-M1 cells) (Fig. 1B,D). To ensure generalized mechanisms of resistance did not underlie emergence of TK216-resistant clones, we tested unrelated anti-cancer toxins etoposide and MLN4924 and confirmed that the clones were specifically resistant to TK216 (Fig. 1C; Supplementary Fig. 1A).

**Figure 1.**
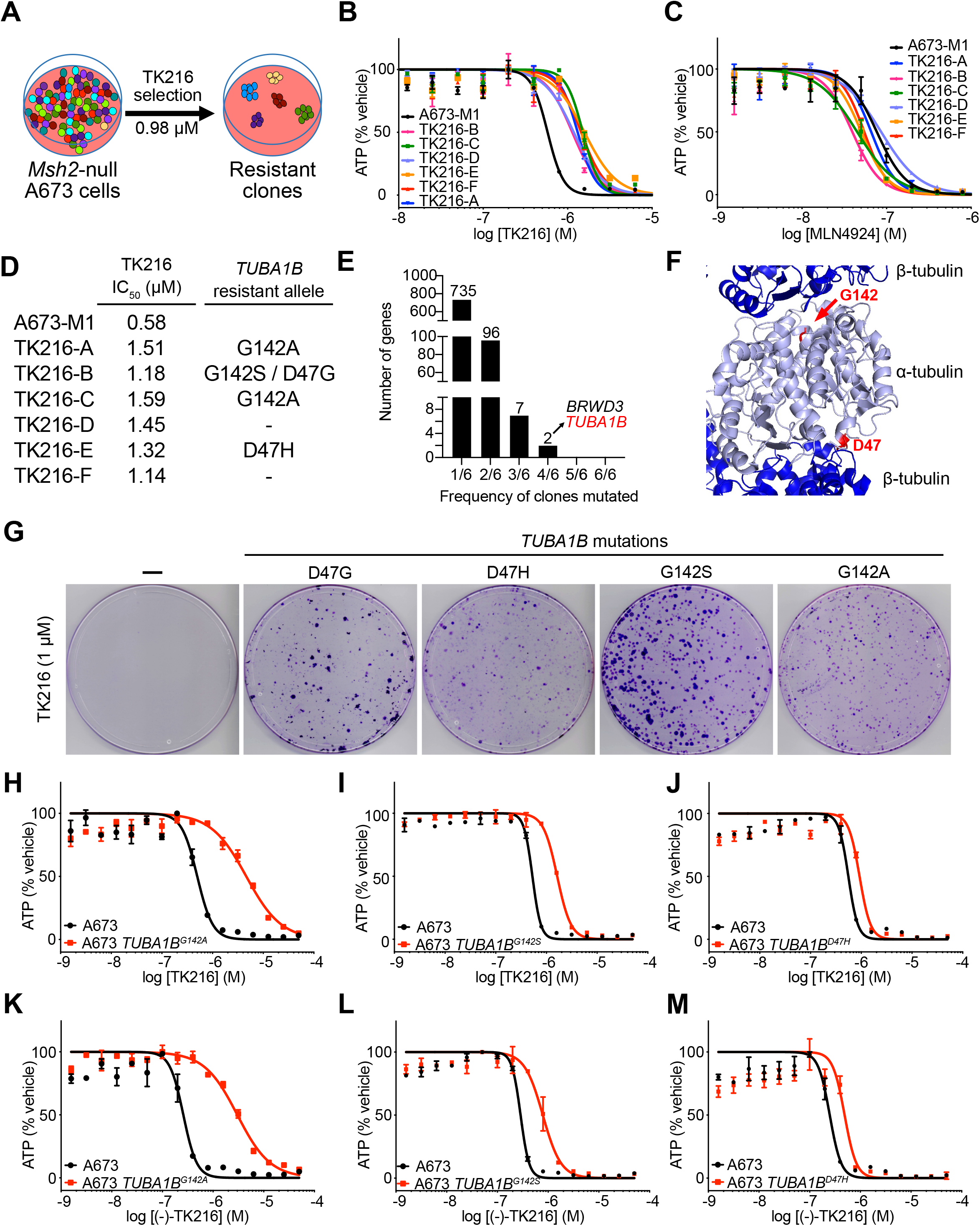
Forward genetic screening for TK216 resistance identifies *TUBA1B* mutations sufficient to confer resistance to YK-4-279/TK216. (A) Workflow for forward genetic screening using Msh2-null EWS cells, A673-M1 cells. Dose-response curves for TK216 (B) and MLN4924 (C) against parental Msh2-null EWS cells, A673-M1, and compound resistant clones. Dose-response curves were performed at least twice for each compound and in duplicate per concentration. (D) Table with half-maximal inhibitory concentrations (IC_50_) for TK216 against parental Msh2-null EWS cells, A673-M1, and compound resistant clones. (E) *TUBA1B* is one of the two genes recurrently mutated in four out of six clones. (F) Crystal structure of *α*-/β-tubulin (PDB: 1Z2B). Highlighted in red are both mutated codons, G142 and D47, found in EWS TK216-resistant cells. (G) Crystal violet staining of EWS cells edited in *TUBA1B* harboring D47G/H or G142S/A mutations. (H-J) Dose-response curves for TK216 against EWS cells harboring *TUBA1B*^*G142A*^, *TUBA1B*^*G142S*^, and *TUBA1B*^*D47H*^ mutations. (K-M) Dose-response curves for (−)-TK216 against EWS cells harboring *TUBA1B*^*G142A*^, *TUBA1B*^*G142S*^, and *TUBA1B*^*D47H*^ mutations. Dose-response curves were performed at least twice for each compound and in duplicate per concentration.

We identified recurrently mutated genes in the TK216-resistant clones by exome sequencing^15^. Two genes, *TUBA1B* and *BRWD3*, were mutated in four out of six clones, with no other gene recurrently mutated in more than four clones (Fig. 1E). No mutations were identified in *EWSR1*, *FLI1*, or *DHX9* (Supplemental Table). We compared somatic mutations between TK216-resistant clones to establish whether clones were related or arose independently. TK216-A and TK216-C clones shared 91 mutations, suggesting that these clones were closely related. No other clones shared more than 11 somatic mutations. We prioritized *TUBA1B*, encoding α-tubulin, as a candidate gene because independent clones harbored recurrent mutations in two codons leading to different amino acid substitutions, G142A/S and D47G/H. The observation of different substitutions of the same codon suggested strong selective pressure for alteration of these specific residues of tubulin in the presence of TK216 (Fig. 1D,F). Interestingly, the G142S mutation was previously reported to confer resistance to the MT destabilizing agent dinitroaniline, raising the possibility that TK216 targeted MTs^16^.

We used CRISPR/Cas9 to engineer each mutation into A673 EWS cells to determine whether codon 47 and codon 142 mutations in *TUBA1B* were sufficient to induce resistance to either YK-4-279 or TK216 (Methods). Cells nucleofected with CRISPR-Cas9 components were selected with TK216 (1 μM) for 2 weeks followed by crystal violet staining. Emerging clones were observed in cells transfected with the *TUBA1B* mutation repair templates whereas no clones were visible in the control condition without Cas9 protein or the repair template (Fig. 1G). This result suggested that *TUBA1B* codon 47 or 142 mutation was sufficient to confer resistance to TK216.

We expanded resistant pools of *TUBA1B*^*G142*^ and *TUBA1B*^*D47*^ cells and validated mutations by Sanger sequencing (Supplementary Fig. 2A-C). The engineered *TUBA1B*^*G142A*^ mutation appeared to be homozygous, whereas the *TUBA1B*^*G142S*^ and *TUBA1B*^*D47H*^ mutations were heterozygous or present in one third of alleles, respectively. We performed dose-response curves of parental and mutant cell pools. *TUBA1B*^*G142S*^, *TUBA1B*^*G142A*^, and *TUBA1B*^*D47H*^ mutations were independently sufficient to confer resistance to YK-4-279 and TK216 (Fig. 1H-J; Supplementary Fig. 2D-F). *TUBA1B* ^*G142A*^ cells exhibited the greatest degree of resistance to TK216, possibly relating to homozygosity of the engineered mutation. A673-*TUBA1B*^*G142A*^, -*TUBA1B*^*G142A*^, and -*TUBA1B*^*D47H*^ cells were not resistant to the DNA polymerase α inhibitor CD437 or the neddylation-activating enzyme inhibitor MLN4924 (Supplementary Fig. 2G-L).

Both YK-4-279 and TK216 contain a chiral center, and previous studies demonstrated that the (−)-YK-4-279 enantiomer was responsible for the anti-cancer activity in EWS cells^17^. We separated (+)-TK216 and (−)-TK216 enantiomers to 98.8% and 99.4% purity, respectively, using supercritical fluid chromatography (SFC) (Lotus Separations, LLC). Consistent with prior reports, (−)-TK216 enantiomer exhibited 56-fold greater anti-cancer activity in Ewing sarcoma cells (IC_50_ = 0.26 μM), compared to the (+)-TK216 enantiomer (IC_50_ = 14.57 μM) (Supplementary Fig. 3A,B). *TUBA1B*^*G142S*^, *TUBA1B*^*G142A*^, and *TUBA1B*^*D47H*^ mutations were also sufficient to confer resistance to purified (−)-TK216 enantiomer (Fig. 1K-M). Thus, introduction of a single *TUBA1B* mutation identified by forward genetics screening in EWS cells was sufficient to endow resistance to YK-4-279, TK216, and (−)-TK216 enantiomer.

Review of the crystal structure of the α-tubulin:β-tubulin dimer placed G142 and D47 mutations at opposite interfaces of the heterodimer, making it unlikely that these mutations impaired interaction of TK216 by altering a single binding pocket (Fig. 1F). We hypothesized that TK216 might act as an MT destabilizing agent, and that G142 and D47 mutations induce resistance to TK216 by stabilizing MTs. We therefore tested whether G142A and G142S mutations also conferred resistance to other MT destabilizing agents. Indeed, A673-*TUBA1B*^*G142A*^, -*TUBA1B*^*G142A*^, and -*TUBA1B*^*D47H*^ cells exhibited resistance to colchicine (Supplementary Fig. 2M-O).

These studies suggested a model in which anti-proliferative activity of YK-4-279 and TK216 stemmed from their action as MT destabilizing agents in EWS cells. We sought to reconstitute MT polymerization *in vitro* to directly assess whether YK-4-279 and TK216 altered MT function. MTs are dynamic structures composed of α-tubulin:β-tubulin heterodimers that polymerize and de-polymerize through a phenomenon called dynamic instability^18^. We used an MT turbidity assay to determine whether YK-4-279 and TK216 impacted MT dynamics. This assay measures the formation of MT polymers by reading absorbance of a mixture of α-tubulin:β-tubulin heterodimers in conditions that facilitate polymerization. Using a molar ratio of compound:tubulin (2:1), 5 μM of YK-4-279 and TK216 inhibited MT polymerization. The positive control, colchicine (5 μM), also potently suppressed MT polymerization (Fig. 2A). Inhibition of MT polymerization was evident with 0.5 μM of TK216 and increased in a dose dependent manner to the maximum concentration tested (20 μM) (Fig. 2B). To exclude nonspecific small molecule assay interference, we tested CD437 and observed no impact on MT polymerization (Fig. 2A).

**Figure 2.**
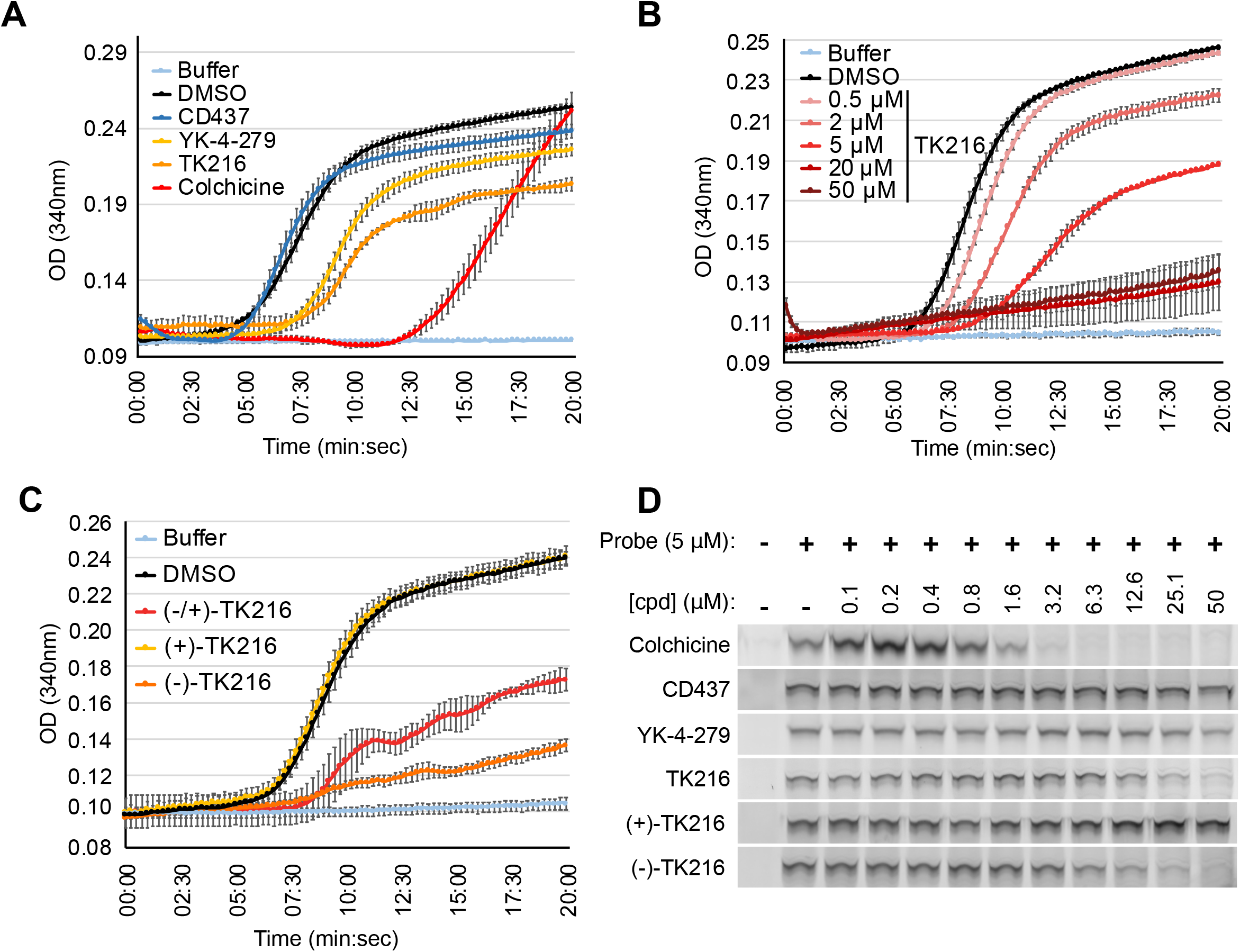
YK-4-279 and TK216 binds at the colchicine site and inhibit MT polymerization *in vitro*. (A) Assessing MT polymerization after addition of DMSO, CD437, YK-4-279, TK216, or colchicine at 5 μM. (B) Dose dependent inhibition of MT polymerization upon TK216 treatment at 0.5 μM, 2 μM, 5 μM, 20 μM, and 50 μM. (C) Inhibition of MT polymerization by (−)-TK216 enantiomer compared to inactive (+)-TK216 enantiomer. All samples were performed in triplicates. (D) Cell-based competition assay using covalent β-tubulin probe to assess tubulin binding by compounds, colchicine, CD437, YK-4-279, TK216, (+)-TK216, and (−)-TK216.

We next evaluated whether inhibition of MT polymerization was enantiomer specific. Indeed, (−)-TK216 potently inhibited MT polymerization whereas (+)-TK216 did not disrupt MT polymerization (Fig. 2C). Therefore, the anti-proliferative activity and inhibition of MT polymerization were both unique properties of the (−)-enantiomer of TK216.

Treatment of cells and xenografts with YK-4-279 was reported to synergize with vincristine; however, the mechanism underlying synergy has not been elucidated. If YK-4-279 and vincristine both target MTs, how can synergy between these agents be explained? Distinct chemical families target MTs through several different binding pockets^19^. We hypothesized that synergy between TK216 and vincristine could be explained if these agents acted on distinct MT binding pockets. We previously reported the development of a tubulin chemical probe that covalently modifies Cys239 within the colchicine binding pocket of β-tubulin^14^. We developed a cell-based competition assay using the tubulin chemical probe and showed that small molecules acting through the colchicine binding pocket competed the benzamide probe whereas vincristine, which acts through a separate vinca alkaloid binding pocket, did not^14^.

To test whether TK216 and vincristine acted through the same or different MT binding pockets, we performed tubulin probe competitions with TK216 in A673 EWS cells (Methods). We confirmed the fidelity of the probe competition assay by testing chemically distinct MT destabilizing agents that act through the colchicine binding pocket, colchicine, rigosertib and tivantinib^20–23^. Indeed, colchicine, rigosertib, and tivantinib competed at concentrations above 0.8, 1.6, and 6.3 μM respectively. In contrast, CD437 exhibited no probe competition at concentrations up to 50 μM, which was well above the IC_50_ for cytotoxicity (∼600 nM) (Fig. 2D; Supplementary Fig. 4).

We next determined whether YK-4-279 and TK216 competed the tubulin chemical probe. Both compounds competed the benzamide probe at concentrations above 12.5 μM (YK-4-279) and 6.3 μM (TK216). We also performed tubulin probe competition assays using the purified TK216 enantiomers (Fig. 2D). (−)-TK216 potently competed the benzamide probe at concentrations above 1.6 μM, whereas (+)-TK216 was devoid of tubulin probe competition activity at concentrations up to 50 μM (Fig. 2D; Supplementary Fig. 5). Therefore, YK-4-279/TK216 and vincristine destabilize MTs through distinct binding mechanism. These results also demonstrate that binding to MTs in cells, inhibition of MT polymerization in vitro, and cytotoxicity (see Fig S1) were unique properties of the (−)-TK216 enantiomer.

Each experimental approach presented here, including unbiased forward genetics, reconstituted MT polymerization, and cell-based chemical probe assays converged upon the singular conclusion that TK216 exhibits anti-proliferative activity by acting as an MT destabilizing agent. Can these findings be reconciled with the scientific literature suggesting that YK-4-279 and TK216 act directly on the EWSR1-FLI1 fusion protein? YK-4-279 was identified using a surface plasmon resonance assay to identify small molecules capable of binding recombinant EWSR1-FLI1 protein^2^. At high concentration *in vitro* (30 μM), YK-4-279 was found to displace binding of a 10 amino acid peptide from RNA helicase A to recombinant EWSR1-FLI1. The authors also reported that YK-4-279 blocked immunoprecipitation of RNA helicase A and EWSR1-FLI1 in EWS cells and suppressed EWSR1-FLI1-dependent transcription of the *NR0B1* promoter, a validated EWSR1-FLI1 transcriptional target^24^.

Treatment of EWS cells with YK-4-279 induced cell cycle arrest at the G_2_-M transition, a hallmark phenotype of MT agents. In contrast, genetic suppression of EWSR1-FLI1 induced cell cycle arrest at a different checkpoint: the G_1_-S transition^10^. In addition, after the publication of Erkizan et al., a series of publications from independent laboratories demonstrated that YK-4-279 induced cell death in several different cancer types, including prostate cancer, neuroblastoma, lymphoma, melanoma and thyroid cancer^5–9^. The discordance in the phenotypes induced by YK-4-279 and genetic suppression of EWSR1-FLI1, the induction of G_2_-M cell cycle arrest, and the broad anti-proliferative activity of YK-4-279 are consistent with the data presented here and the conclusion that YK-4-279 acts as an MT destabilizing agent. Whether the molecule also exhibits anti-cancer activity through binding EWSR1-FLI1 will require additional medicinal chemistry efforts to decouple such activity from MT destabilization.

Based on our chemical probe competition assays, we favor the hypothesis that TK216 acts on tubulin through the colchicine binding site. However, it is important to recognize the possibility that TK216 binds outside the colchicine pocket and displaces the benzamide probe through an allosteric, or indirect, mechanism. It was also of interest that we identified recurrent mutations impacting α-tubulin, rather than β-tubulin in TK216-resistant A673 clones. The colchicine binding pocket is encoded within β-tubulin, not α-tubulin^25^. If TK216 competes our colchicine binding pocket chemical probe, then why were mutations in β-tubulin not identified? β-tubulin and α-tubulin are each expressed as multiple isoforms from eight genes. Expression of the different tubulin genes varies between cell lineages and cancer cell lines, and many cell lines express relatively uniform levels of multiple β- and α- tubulin genes. Therefore, a compound resistant allele in a single tubulin gene is expected to result in limited impact on MT dynamics or resistance to an MT agent. Interestingly, whereas multiple β-tubulin isoforms are expressed at similar levels in A673 cells, *TUBA1B* is the dominantly expressed α-tubulin (data not shown). We hypothesize that the emergence of *TUBA1B* mutations, rather than β-tubulin gene mutations, in our forward genetics screening stems from greater impact of a single mutation in *TUBA1B* on the cellular pool of MTs.

This study raises the possibility that agents targeting MTs through the colchicine binding pocket might offer clinical benefit in EWS and other tumors. Treatment with TK216 and vincristine induced tumor regression in a subset of patients enrolled in the phase I/II trial^4^. Our observation that TK216 and vincristine act through different binding sites provides a mechanistic explanation for the clinical activity of this combination. Rigosertib, which was initially developed as a Polo-like kinase 1 (PLK1) inhibitor, was subsequently shown to act as an MT destabilizing agent through the colchicine binding pocket^20–23^. Rigosertib, which has advanced to phase III clinical trials, likely represent the first colchicine binding pocket agent to exhibit a reasonable safety profile. The experience with rigosertib, and now TK216, highlights the importance of carefully uncovering the mechanism of action of small molecule therapeutics developed from target-based in vitro screening. However, the activity of these compounds in clinical trials also hints that MT agents acting through the colchicine binding pocket might offer clinical benefit in EWS and other cancers, alone or in combination with MT agents acting through separate binding pockets such as vincristine.

## Supporting information

Supplemental Table_TK216-resistant clones_Somatic Mutations

HPLC Purity data of YK-4-279, TK216 and enantiomers

## Acknowledgements

This work was supported by grants from the Welch Foundation (I-2040, D.G.M. and I-1422, J.K.D.B.), the National Cancer Institute of the NIH (U54CA231649, D.G.M.), a Disease-Oriented Scholar Award from UT Southwestern Medical Center (D.G.M.), a Clinical Investigator Award from the Damon Runyon Cancer Research Foundation (102-19, D.G.M.), the Cancer Prevention and Research Institute of Texas (RP190141, D.G.M.), and a Pilot Synergy Award from the UT Southwestern Dean’s Circle of Friends (D.G.M. and J.K.D.B.). J.K.D.B. holds the Julie and Louis Beecherl, Jr., Chair in Medical Science. We thank Deepak Nijhawan for critically reading the manuscript and innumerable scientific discussions.

## Author contributions

D.G.M. conceived the study. D.G.M. supervised research. J.M.P. designed and performed experiments. K.L., V.L., and X.B. performed experiments. R.R. and J.K.D.B. designed and synthesized chemical probes. J.K. analyzed exome sequencing data. Y.X. supervised J.K. and exome sequencing analysis. D.G.M. and J.M.P. wrote the manuscript.

## METHODS

### Cell Lines

Ewing sarcoma A673 cell lines were cultured at 37°C and 5% CO_2_ in RPMI (R8758, Sigma-Aldrich) and supplemented with 10% FBS (#35-150-CV, Corning), 2 mM L-glutamine (G7513, Sigma-Aldrich), and penicillin/streptomycin (P0781, Sigma-Aldrich). Cells were pasaged using trypsin (T4049, Sigma-Aldrich) every 3-4 days. Parental A673 cell lines are derived from a female subject and were authenticated by STR profiling. Mouse small cell lung cancer 518T2 cells were previously reported ^14^. These cells were cultured in DMEM (D6429, Sigma-Aldrich) supplemented with 5% FBS (#35-150-CV, Corning), 2 mM L-glutamine (G7513, Sigma-Aldrich), and penicillin/streptomycin (P0781, Sigma-Aldrich). Cells were passaged using trypsin (T4049, Sigma-Aldrich) every 3-4 days.

### Compounds

Small molecule CP35 was purchased from ChemBridge (#7658470). Small molecule CP68 was purchased from ChemDiv (#5353-0933). Etoposide was purchased from Sigma-Aldrich (#E1383-100MG). CD437 was purchased from Sigma-Aldrich (#C5865). MLN4924 was purchased from ApexBio (#B1036). Rigosertib was purchased from Cayman Chemical (#15553). Colchicine was purchased from Sigma-Aldrich (#C9754-100MG). YK-4-279 was purchased from Selleck Chemicals (#S7679). TK216 was purchased from MedChem Express (#HY-122903). TK216 enantiomers were separated and purified by Lotus Separations. SFC (supercritical fluid chromatography) separation of 29 mg racemic TK216 yielded 14 mg of (+)-TK216 and 14 mg of (−)-TK216. The separation method used was: AS-H (2 × 25 cm), 35% ethanol/CO_2_ (100 bar), 50 mL/min, 220 nm. Injected volume 1 mL, 2 mg/mL methanol:DCM. YK-4-279, TK216, and both TK216 enantiomers exhibited between 98 and 99% purity as determined by LC-MS analysis performed on an Agilent 1290 HPLC system using an Eclipse XDB-C18 column (46 × 150 mm, 5 μm); Agilent) that was coupled to an Agilent 6130 mass spectrometer run in ESI mode in both positive and negative ionization with a scan range of 100-1,100 m/z. Liquid chromatography was carried out at a flow rate of 0.5 mL/min at 20 °C with a 5 μL injection volume, using the gradient elution with aqueous acetonitrile containing 0.1% formic acid. The gradient was adjusted based on the different polarity of different compounds. All compounds were diluted in DMSO (Sigma-Aldrich, D650-100ML). HPLC chromatograms for all compounds used in the study shown in Figure S5.

### Forward Genetic Screen of TK216

Previously described mismatch-repair deficient EWS cells, A673-M1, were utilized. We first identified the concentration of TK216 that killed 100% of MMR-deficient and -proficient A673 cells after 1 week of compound exposure (IC_100_^1wk^). IC_100_ ^1wk^ determination for TK216 was performed in a 12-well plate seeding 25,000 cells per well. After 24h, TK216 was dispensed using TECAN D300e setting up a minimum concentration of IC_50_^72h^ and a maximum concentration of IC_100_^72h^. Media and TK216 were replenished after 3-4 days. After 7 days, cell viability was determined visually to determine IC_100_ ^1wk^. Then, A673-M1 cells and A673 parental cells were plated in 5 × 10cm plates for each cell line (1 million cells per plate). The following day, TK216 was added at 5 different concentrations: IC_100_^1wk^ ÷ 1.5 (0.66 μM), IC_100_^1wk^ ÷ 1.25 (0.78 μM), IC_100_^1wk^ (0.98 μM), IC_100_^1wk^ × 1.25 (1.22 μM), and IC_100_^1wk^ × 1.5 (1.48 μM) to the plates. Media with TK216 was replenished every 3 – 4 days over the course of 2 weeks. Surviving clones were expanded.

### Whole Exome Sequencing Analysis

Trim Galore (https://www.bioinformatics.babraham.ac.uk/projects/trim_galore/) was used for quality and adapter trimming. The human reference genome sequence and gene annotation data, hg38, were downloaded from Illumina iGenomes (https://support.illumina.com/sequencing/sequencing_software/igenome.html). The sequencing reads were aligned to the genome sequence using Burrows-Wheeler Aligner (BWA, v0.7.17) ^26^. Picard (2.21.3) (https://broadinstitute.github.io/picard) was used to remove PCR duplicates and Genome Analysis Toolkit (GATK, 4.1.4.0) ^27,28^ was used to recalibrate base qualities. Calling variants and genotyping were performed using GATK HaplotypeCaller and the variant calls were filtered by applying the following criteria: QD (Variant Confidence/Quality by Depth) < 2, FS (Phred-scaled p-value using Fisher’s exact test to detect strand bias) > 60, MQ (RMS Mapping Quality) < 40, DP (Approximate read depth) < 3, GQ (Genotype Quality) < 7. Custom Perl scripts (https://github.com/jiwoongbio/Annomen) were used to annotate variants with human transcripts, proteins, and variations (RefSeq and dbSNP build 151) and calculate variant allele frequencies. We defined acquired somatic mutations for each A673-M1 TK216-resistant clones by VAF > 0.2 and VAF < 0.01 for the parental A673-M1 cell line and VAF < 0.05 for previously reported MLN-resistant clones ^11^. Non-coding mutations were excluded from the analysis.

### Introduction of *TUBA1B* Mutations

We performed homology-directed repair using Alt-R CRISPR-Cas9 System and ultramer oligo from Integrated DNA Technologies (IDT) in A673 EWS cells. To prepare the gRNA complex for *TUBA1B*^*G142*^ and *TUBA1B*^*D47*^, we combined 10 μL of Alt-R CRISPR-Cas9 crRNA (100 μM) (*TUBA1B*^*G142*^ sequence: 5’-UUC UUG GUU UUC CAC AGC UUG UUU UAG AGC UAU GCU-3’; *TUBA1B*^*D47*^ sequence: 5’-CUC ACU GAA GAA GGU GUU GAG UUU UAG AGC UAU GCU-3’) and 10 μL Alt-R CRISPR-Cas9 tracrRNA (100 μM). The mixture was heated to 95°C for 5 minutes, then allowed to slowly cool to room temperature. Ribonucleoprotein complex with Cas9 was formed by combining 3 μL of gRNA complex with 2 μL Alt-R Cas9 enzyme and incubating at RT for 10-20 minutes. Two million A673 cells were resuspended in 120 μL of SF Cell Line 4D-NucleofectorTM X Kit L (Cat. #: V4XC-3024). The transfection mix was prepared using: 15 μL of RNP complex, 3.6 μL of 100 μM Ultramer ssODN donor (*TUBA1B*^*G142S*^ sequence: 5’-A*C*C AGT GCA CCG GTC TTC AGG GCT TCT TGG TTT TCC ACA GCT TTA GTG GGG GAA CTG GTT CTG GGT TCA CCT CCC TGC TCA TG*G*A-3’; *TUBA1B*^*G142A*^ sequence: 5’-A*C*C AGT GCA CCG GTC TTC AGG GCT TCT TGG TTT TCC ACA GCT TTG CTG GGG GAA CTG GTT CTG GGT TCA CCT CCC TGC TCA TG G*A*A-3’; *TUBA1B*^*D47H*^ sequence: T*G*G CCA GAT GCC AAG TGA CAA GAC CAT TGG GGG AGG AGA TGC CTC CTT CAA CAC ATT CTT CAG TGA GAC GGG CGC TGG CAA GCA CGT GCC CCG GGC T*G*T; *TUBA1B*^*D47G*^ sequence: T*G*G CCA GAT GCC AAG TGA CAA GAC CAT TGG GGG AGG AGA TCA CTC CTT CAA CAC ATT CTT CAG TGA GAC GGG CGC TGG CAA GCA CGT GCC CCG GGC T*G*T) and added to 60 μL of previously prepared cell suspension. Nucleofection was performed using 4D-Nucleofector™ core unit from LONZA. Cells were plated into one well of a 6-w plate after nucleofection in 2 mL of RPMI (10% FBS). After one week, sham cells or ssODN (*TUBA1B*^*G142S/A*^) cells were plated into 10 cm dishes. Each sample was plated into three 10cm dishes using 1 million cells per dish. *TUBA1B*^*G142S/A*^ cells were treated with TK216 at concentrations: 0.75 / 1 / 1.25 μM. Media and small molecules were replenished every 3 – 4 days over the course of 2 weeks followed by expansion in media without compound for 1 week. A673-*TUBA1B*^*G142S/A*^ and -*TUBA1B*^*D47H/G*^ cells were stained with crystal violet staining solution prepared with 1% (weight/volume ratio) crystal violet from Sigma-Aldrich (#C6158-50G) in 10% ethanol. After TK216 selection, one dish with A673-*TUBA1B*^*G142S*^, -*TUBA1B*^*G142A*^, and -*TUBA1B*^*D47H*^ cells were expanded to performed DRC to validate compound resistance.

### Cytotoxic Assay

For mouse SCLC, 518T2, cells were seeded in duplicates in 96-well plates, 10,000 cells and 200 μL of DMEM media (5% FBS) per well. For human EWS, A673, cells were seeded in duplicate in 96-well plates, 3,000 cells and 200 μL of RPMI media (10% FBS) per well. After overnight incubation, compounds were dispensed using a D300e Digital Dispenser (TECAN) in 15-point dose response manner using a maximum and minimum concentration of 50 μM and 1.58 nM, respectively. Cell viability was assessed after 72 hours using CellTiter-Glo luminescent cell viability assay (Promega, #G7571). The CellTiter-Glo reagent was diluted by adding PBS-Triton-X (1%) (1:1 ratio). Each value was normalized to cells treated with DMSO, and the IC50 values were calculated using GraphPad Prism software.

### MT Polymerization Assay

We used cycled-tubulin purchased from PurSolutions (Cat. #: 032005). MT polymerization occurs spontaneously upon incubation of cycled-tubulin in PEM buffer with GTP at 37 Celsius. Each condition was performed in triplicate from a 384-well plate. First, the 384-well plate was prewarmed at 37 Celsius using plate reader Synergy2 (Biotek). Master mix containing cycled-tubulin, PEM buffer and GTP was prepared for the samples analyzed in an Eppendorf tube: 2.5 μg of cycled-tubulin (20 mg/mL), (5X) PEM buffer (400 mM PIPES, 5 mM EGTA, 5 mM MgCl_2_, pH = 6.8), DTT (1 mM), GTP (1 mM), DMSO (3%), and ddH_2_O up to 30 μL per reaction and incubateed on ice 3-5 minutes. After incubation, 30 μL of the master mix was added per well in a 384-well plate. Immediately after compounds were added at desired concentration using a D300e Digital Dispenser (TECAN). Absorbance at 340 nm was measured immediately after every 15 seconds for 20 minutes.

### Cell-based Tubulin Competition Assay

Murine SCLC cells, 518T2, were seeded in 12-w plates (1 million cells per well) with 1 mL DMEM media (5% FBS) each well. Human EWS cells, A673, were seeded in 12-w plates (0.5 million cells per well) with 1 mL RPMI media (10% FBS) each well. Small molecules were dispensed 24-hours later at increasing concentrations using a D300e Digital Dispenser (TECAN) in 10-point dose response manner using a maximum and minimum concentration of 50 μM and 100 nM. Following 30 minutes incubation at 37 °C in 5% CO_2_, tubulin covalent probe was dispensed at 5 μM in 11 wells. After 30 minutes incubation, cells were washed gently with PBS and then lysed in 1% SDS Buffer A (50 mM HEPES pH 7.4, 10 mM KCl, 2 mM MgCl_2_), freshly supplemented with 1:10,000 benzonase (Sigma-Aldrich). After incubation, copper-mediated click chemistry with a fluorescent azide was performed. Covalently modified β-tubulin was visualized by SDS-PAGE and scanning gels for fluorescence^14^.

## Data availability

WES data for samples TK216 clones are accessible at SRA accession number: PRJNA770630

## OLIGONUCLEOTIDE TABLE

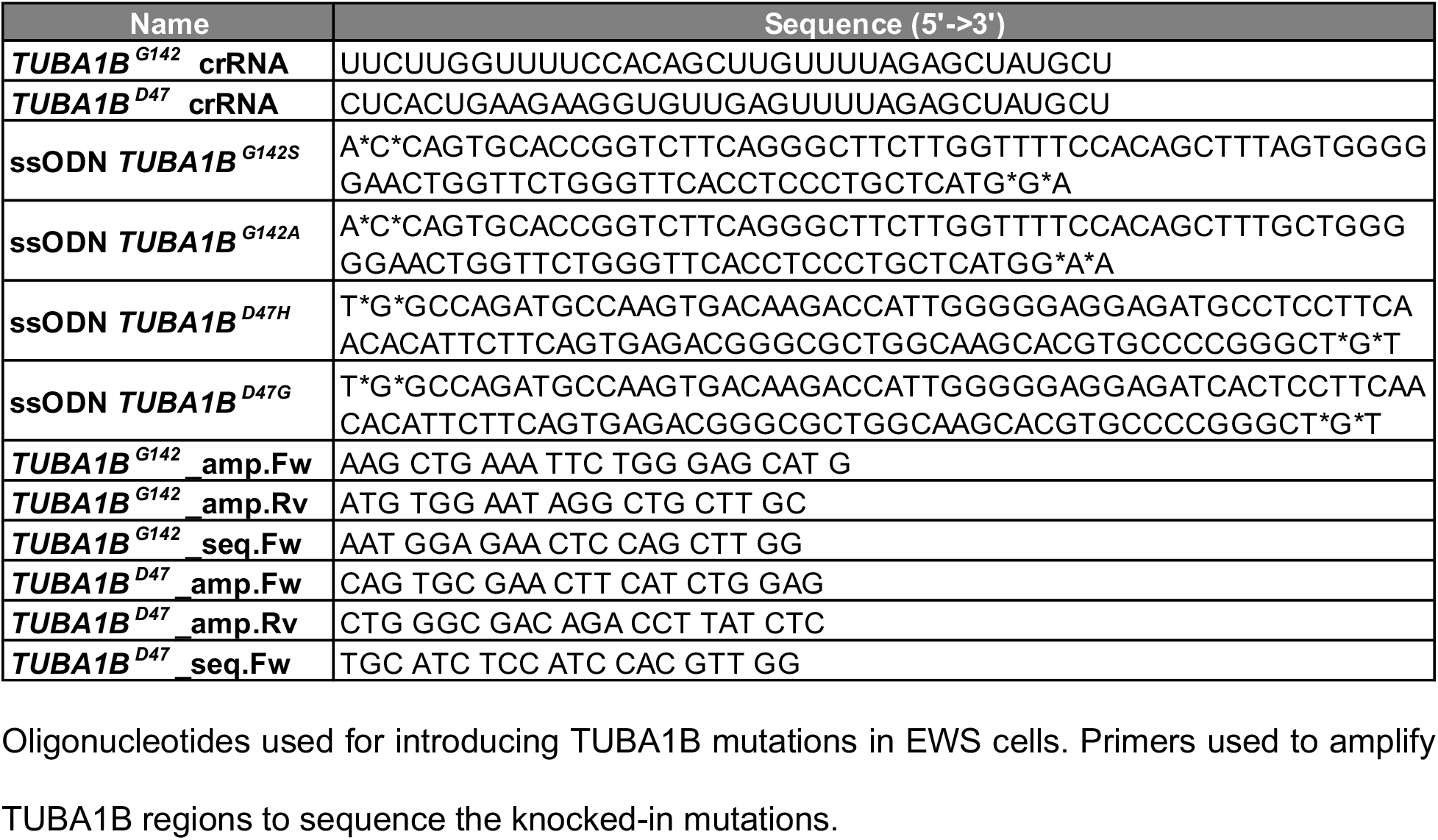

---

**Supplementary Figure 1.**
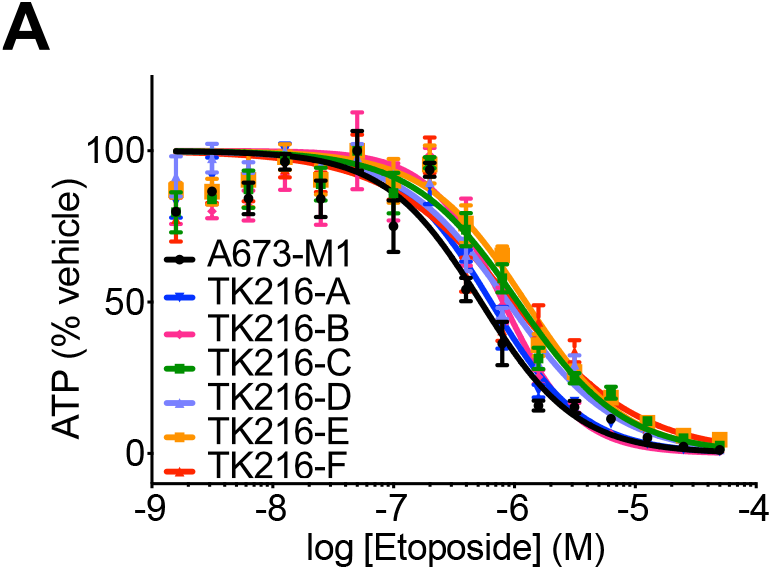
(A) Dose-response curve for etoposide against A673-M1, Msh2-null EWS cells, and TK216 resistant clones.

**Supplementary Figure 2.**
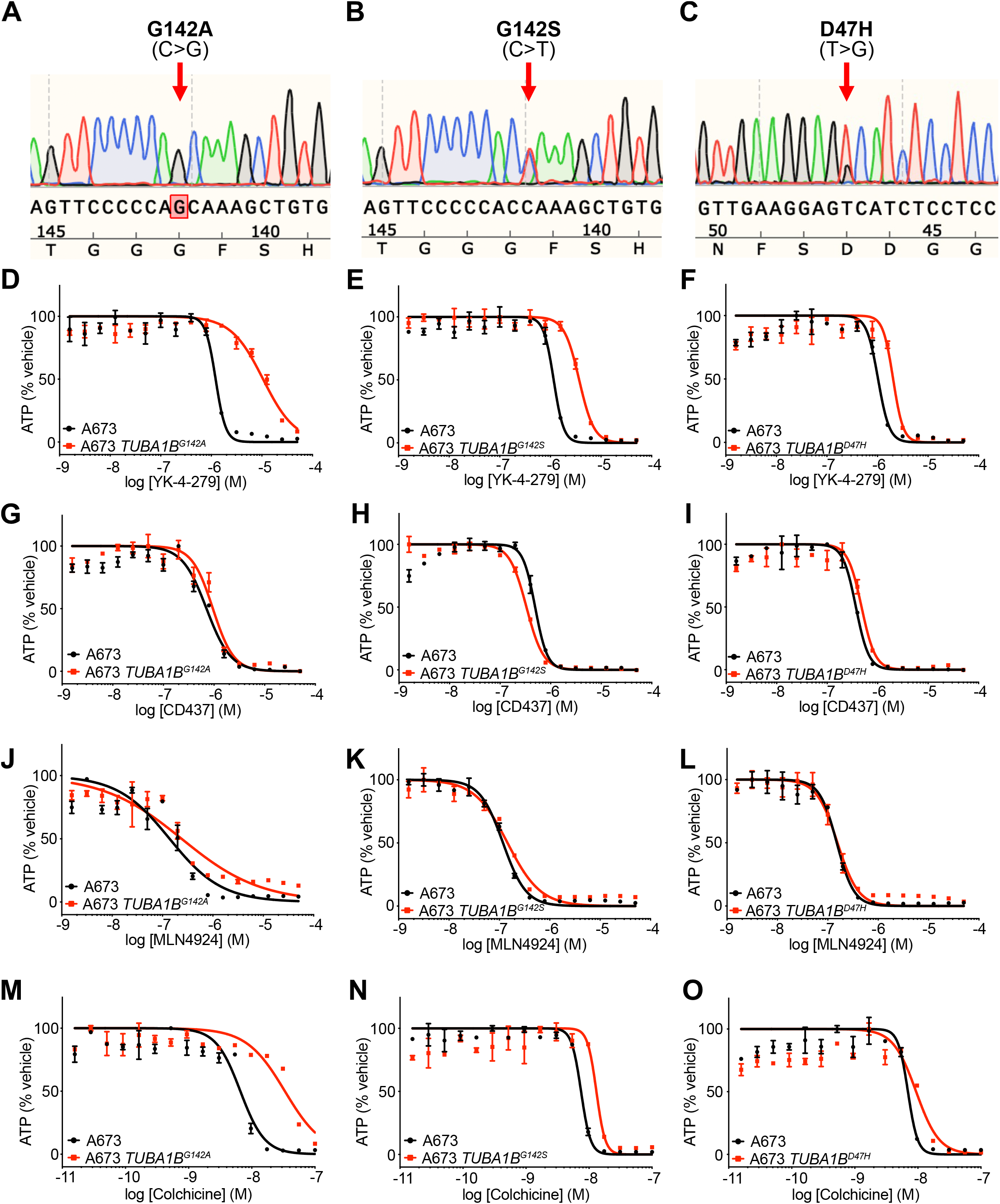
(A,B,C) Sequencing traces for TUBA1B^G142A^, TUBA1B^G142S^, and TUBA1B^D47H^ mutations in EWS cells. (D,G,J,M) Dose-response curves for, YK-4-279 (D), CD437 (G), MLN4924 (J), colchicine (M) against EWS cells harboring TUBA1B^G142A^ mutation. (E,H,K,N) Dose-response curves for, YK-4-279 (E), CD437 (H), MLN4924 (K), colchicine (N) against EWS cells harboring TUBA1B^G142A^ mutation. (F,I,L,O) Dose-response curves for, YK-4-279 (F), CD437 (I), MLN4924 (L), colchicine (O) against EWS cells harboring TUBA1B^D47H^ mutation.

**Supplementary Figure 3.**
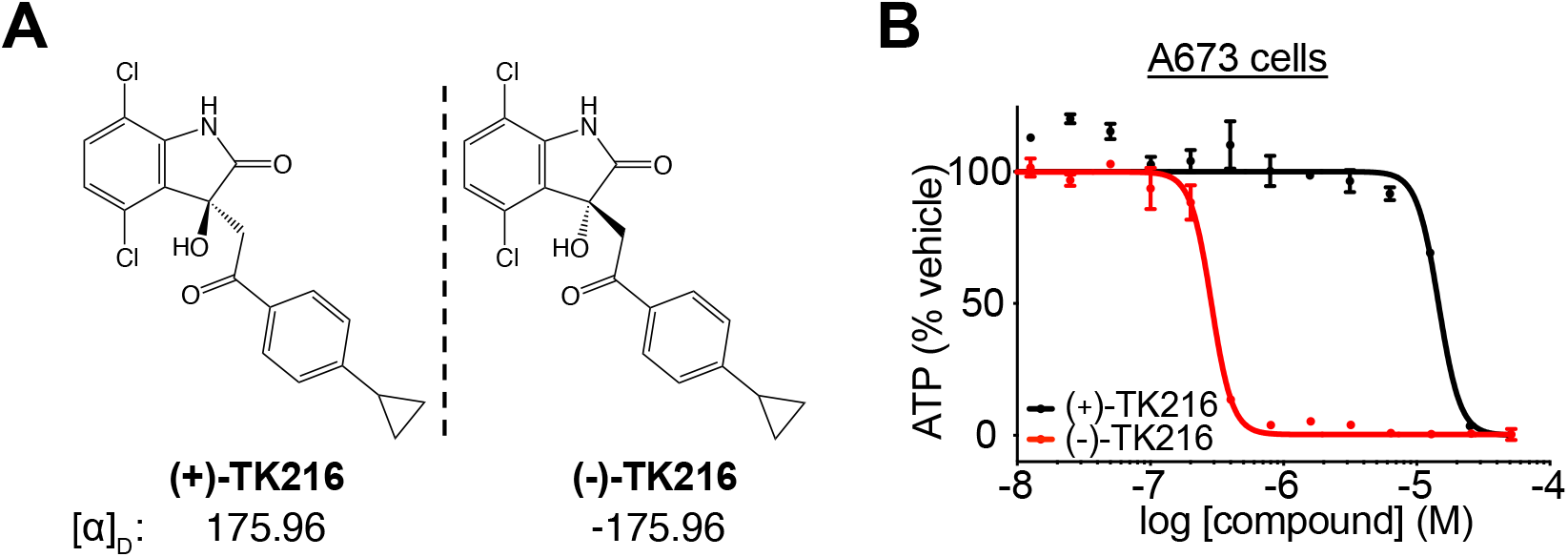
(A) Chemical structure and specific rotation ([α]_D_) of each TK216 enantiomer. (B) Dose-response curve for (−)-TK216 and (+)-TK216, against EWS A673 cells. Densitometry graphs under gels were generated using ImageJ to measure intensity of β-tubulin band. Densitometry data were normalized to the control no competitor plus probe lane.

**Supplementary Figure 4.**
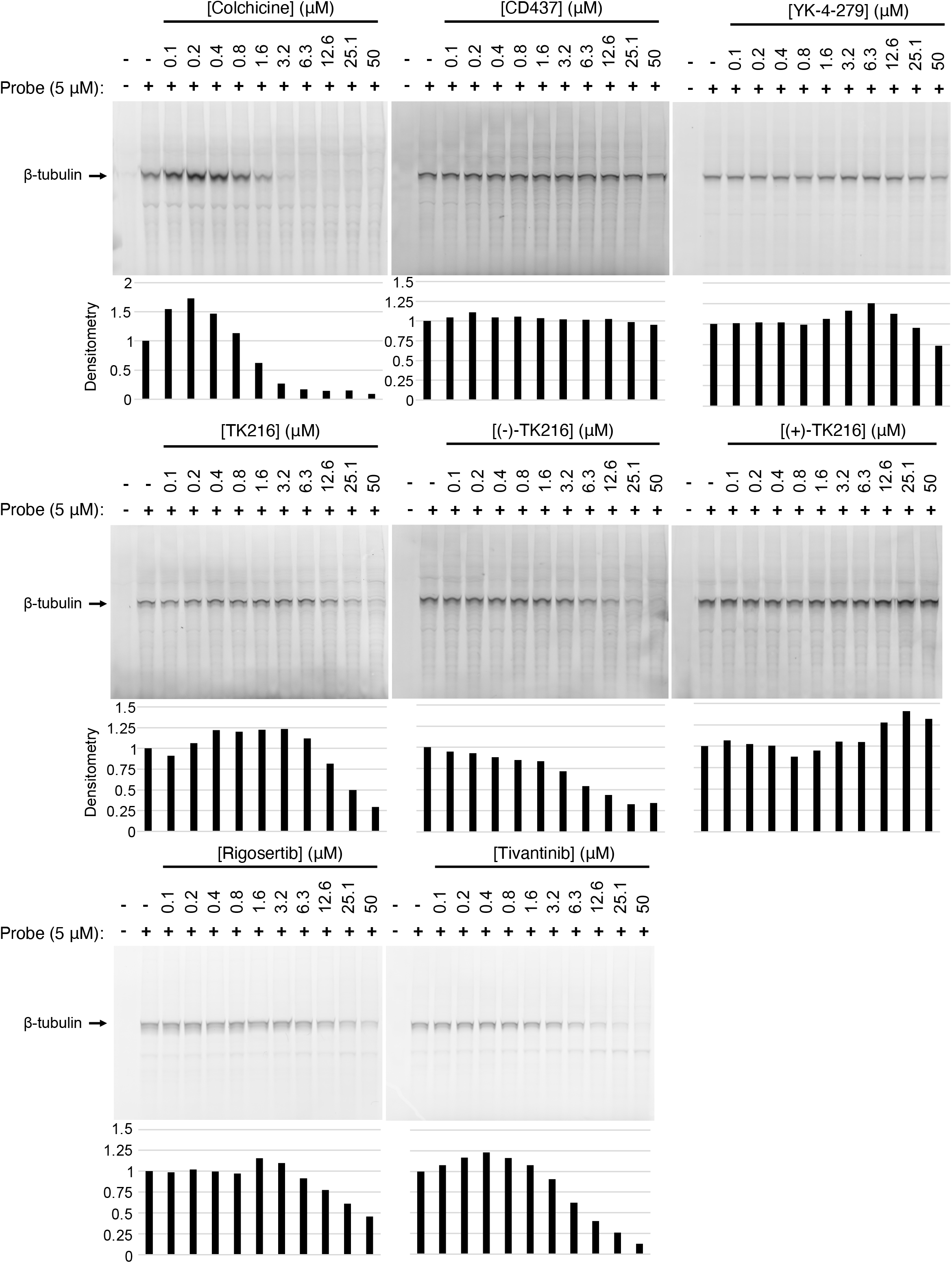
Full gel images of cell-based competition assay using covalent β-tubulin probe to assess tubulin binding in EWS cells by compounds, colchicine, CD437, YK-4-279, TK216, (−)-TK216, (+)-TK216, rigosertib, and tivantinib. Densitometry graphs under gels were generated using ImageJ to measure intensity of β-tubulin band. Densitometry data were normalized to the control no competitor plus probe lane.

