## Supplementary material for "TK216 targets microtubules in Ewing sarcoma cells": HPLC Purity data of YK-4-279, TK216 and enantiomers

A

B

C

D

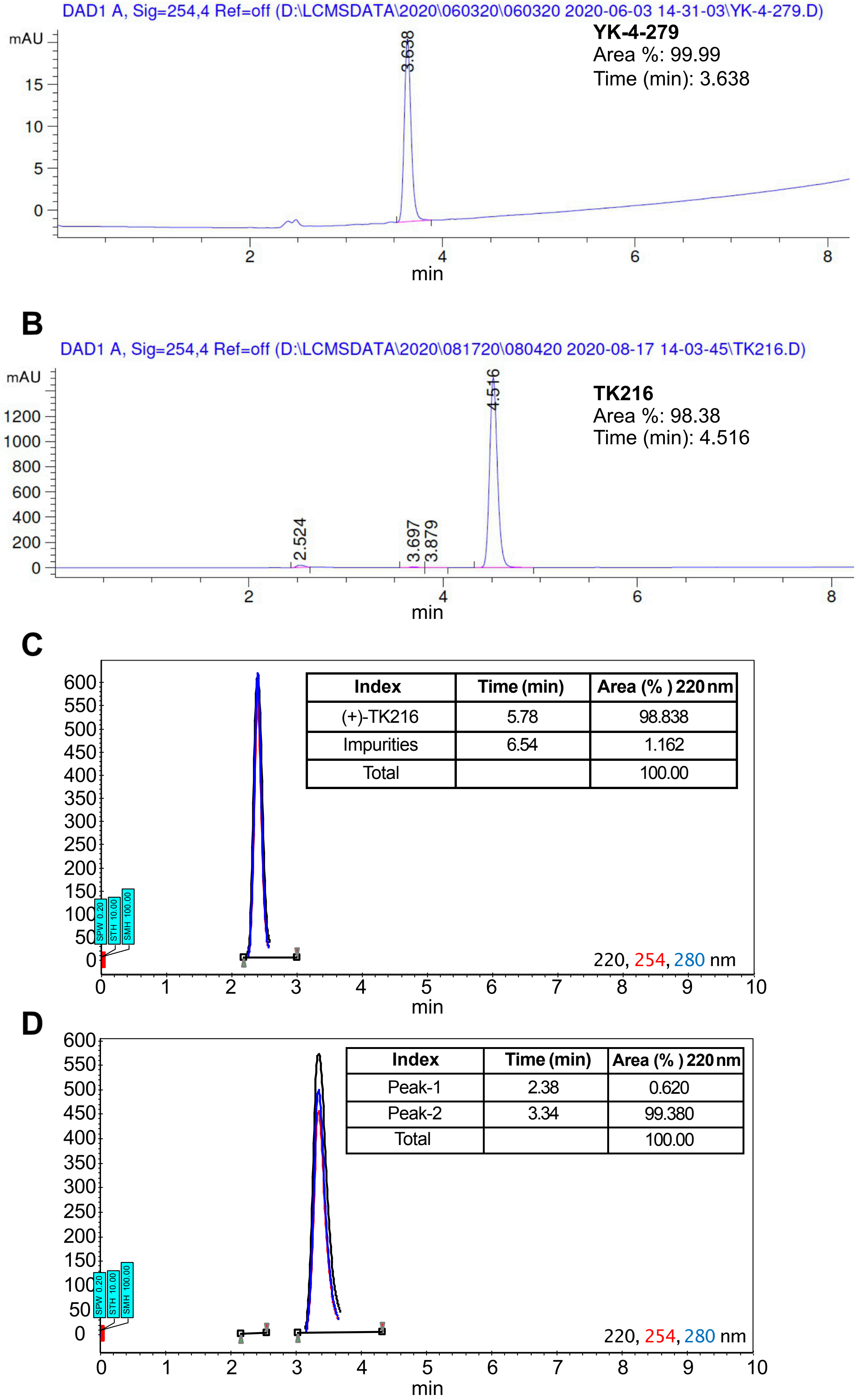


| Index | Time (min) | Area (% ) 220 nm |
|---|---|---|
| (+)-TK216 | 5.78 | 98.838 |
| Impurities | 6.54 | 1.162 |
| Total | | 100.00 |

| Index | Time (min) | Area (% ) 220 nm |
|---|---|---|
| Peak-1 | 2.38 | 0.620 |
| Peak-2 | 3.34 | 99.380 |
| Total | | 100.00 |

**HPLC Chromatograms of YK-4-279 (A), TK216 (B), (+)-TK216 (C), and (-)-TK216 (D).**
